# FORKS: Finding Orderings Robustly using k-means and Steiner trees

**DOI:** 10.1101/132811

**Authors:** Mayank Sharma, Huipeng Li, Debarka Sengupta, Shyam Prabhakar, Jayadeva

## Abstract

Recent advances in single cell RNA-seq technologies have provided researchers with unprecedented details of transcriptomic variation across individual cells. However, it has not been straightforward to infer differentiation trajectories from such data, due to the parameter-sensitivity of existing methods. Here, we present Finding Orderings Robustly using k-means and Steiner trees (FORKS), an algorithm that pseudo-temporally orders cells and thereby infers bifurcating state trajectories. FORKS, which is a generic method, can be applied to both single-cell and bulk differentiation data. It is a semi-supervised approach, in that it requires the user to specify the starting point of the time course. We systematically benchmarked FORKS and eight other pseudo-time estimation algorithms on six benchmark datasets, and found it to be more accurate, more reproducible, and more memory-efficient than existing methods for pseudo-temporal ordering. Another major advantage of our approach is its robustness – FORKS can be used with default parameter settings on a wide range of datasets.

## 1 Introduction

Single cell RNA-seq (scRNA-seq) is a novel technique that allows measurement of transcriptomes at single cell resolution [1, 2, 3, 4, 5, 6, 7, 8]. Such a measurement is a snapshot of ongoing cellular processes. In contrast, bulk-sample methods only provide population-averaged measurements of gene expression, and therefore fail to accurately reflect the underlying diversity of expression phenotypes in the presence of cellular heterogeneity. For example, only single cell analysis can differentiate between high expression in a subset of cells and moderate expression in all cells. Thus, scRNA-seq is extremely valuable in unraveling complex biological phenomena such as cell differentiation, gene expression “noise” and gene regulatory interactions [9]. In particular, one common analytical approach has been to assume that transcriptomic variation across cells within a population approximates variation across time within a single lineage [10, 11, 12].

The problem of ordering single cell transcriptomes along continuous trajectories in state (expression) space, known as pseudo-temporal ordering, is typically solved in a lower-dimensional space that approximates proximity relationships between cells in the full space of transcript abundances [13]. Multiple methods have been used for dimension reduction in this context, including Principal Component Analysis (PCA) [14], Independent Component Analysis (ICA) [15], Diffusion Maps [16] and t-Stochastic Neighborhood Embedding (t-SNE) [17] are popular choices among others.

Multiple tools have been developed for pseudotemporal ordering, some of which are capable of discovering branched trajectories. The initial release of Monocle [18] used ICA as dimensionality reduction technique. Their latest release Monocle2 uses the Discriminative Dimensionalty Reduction Tree (DDRTree) [19] technique to reduce dimensionality. DDRTree solves the dual objective of finding the linear projection such that clusters are well separated. Wishbone [20] uses Diffusion Maps to create a nearest neighbor graph whereas Diffusion Pseudo-time (DPT) [21, 22] uses Diffusion Maps to find the transition probabilities and hence order the cells based on the diffusion distance, Waterfall [23] uses PCA, DPT uses Diffusion Maps, GPfates [24] uses Gaussian process Latent variable model (GPLVM) to reduce the dimension and overlapping mixture of gaussian processes (OMGP) to identify the bifurcations, TSCAN [25] uses a combination of PCA to reduce the dimension and model based clustering to find the cluster centers, SCUBA [26] models the developmental trajectory using a stochastic dynamical model and works in original space or in some cases reduce the dimension using t-SNE. DeLorean [27] uses Gaussian process to learn the gene expression profiles and the pseudo-time associated with each cell. SLICER [28] first select the genes then computes alpha hull of the data to find the optimal nearest neighbors for Locally Linear Embedding (LLE) [29]. Embeddr [30] reduces the dimension of data using Laplacian Eigenmaps [31], then fits principal curves to the manifold and projects onto the curve to estimate the pseudo-time. Embeddr cannot faithfully recover bifurcating trajectories if present in the data. Mpath [32] generates a branched trajectory by joining the cluster centers found using Hierarchial clustering and then projecting onto the Minimum Spanning Tree (MST).

Cannondt et. al. [33], provides a broad overview of many existing pseudo-time estimation algorithms which are compared in terms of qualitative features. However, quantitative benchmarking of pseudo-temporal ordering methods on multiple datasets has been limited. In this study, we therefore evaluated the pseudo-time estimation accuracy of FORKS along with eight other algorithms on three scRNA-seq datasets and two bulk expression profiling datasets for which the true temporal stages are known. We found one major limitation of the existing methods that: the default parameter settings were not robust. As a consequence, they required manual tuning of multiple hyper-parameters for each new dataset. We also found that some of the algorithms designed to detect branching trajectories performed poorly when the underlying biological process was unbranched. Moreover, non-linear embeddings were not necessary for inferring differentiation trajectories. In fact, linear methods such as PCA yielded results comparable to or better than those of non-linear methods.

In order to address the above challenges in pseudo-temporal ordering, we developed FORKS, a method that infers bifurcating trajectories when present, and linear trajectories otherwise without hyper-parameter tuning. FORKS relies on generalization of MST known as Steiner Trees [34] to create robust bifurcating trajectories. As previously noted [25], the MST of the entire set of cells may not be robust, since it could be substantially altered by a small changes in the data. FORKS therefore reduces the complexity of the problem and ameliorates measurement noise by finding the Steiner Points which can be thought of as cluster centers in the case of k-means but connected via an MST. Another advantage of this approach is that FORKS is scalable to thousands of cells and genes. In order to compare the performance of FORKS against existing methods, we performed the first systematic bench-marking exercise involving 9 algorithms in total and 5 transcriptomic datasets for which the true time stamps known.

## 2 Results

### 2.1 Datasets

We use 6 different datasets in the paper namely Arabidopsis [43], Deng 2014 [44], Guo 2010 [45], Klein [46], LPS [47] and Preimplant [48]. Among these datasets [43] is microarray dataset, [45] is Reverse transcription polymerase chain reaction (RT-PCR) dataset, and the rest are single cell RNA-seq datasets. These datasets cover a whole spectrum of techniques that have been used in gene expression analysis. The knowledge of observed cell time for each of the cell enables us to benchmark FORKS with other state-of-the-art algorithms that infer the pseudo-time trajectory. The details of datasets are presented in Table 1. To our knowledge this is the first comprehensive benchmarking with such a varied set of algorithms and datasets. The projection of individual datasets on the the first two principal components after the preprocessing step is shown in Figure 1 (1a-1f)

**Figure 1:**
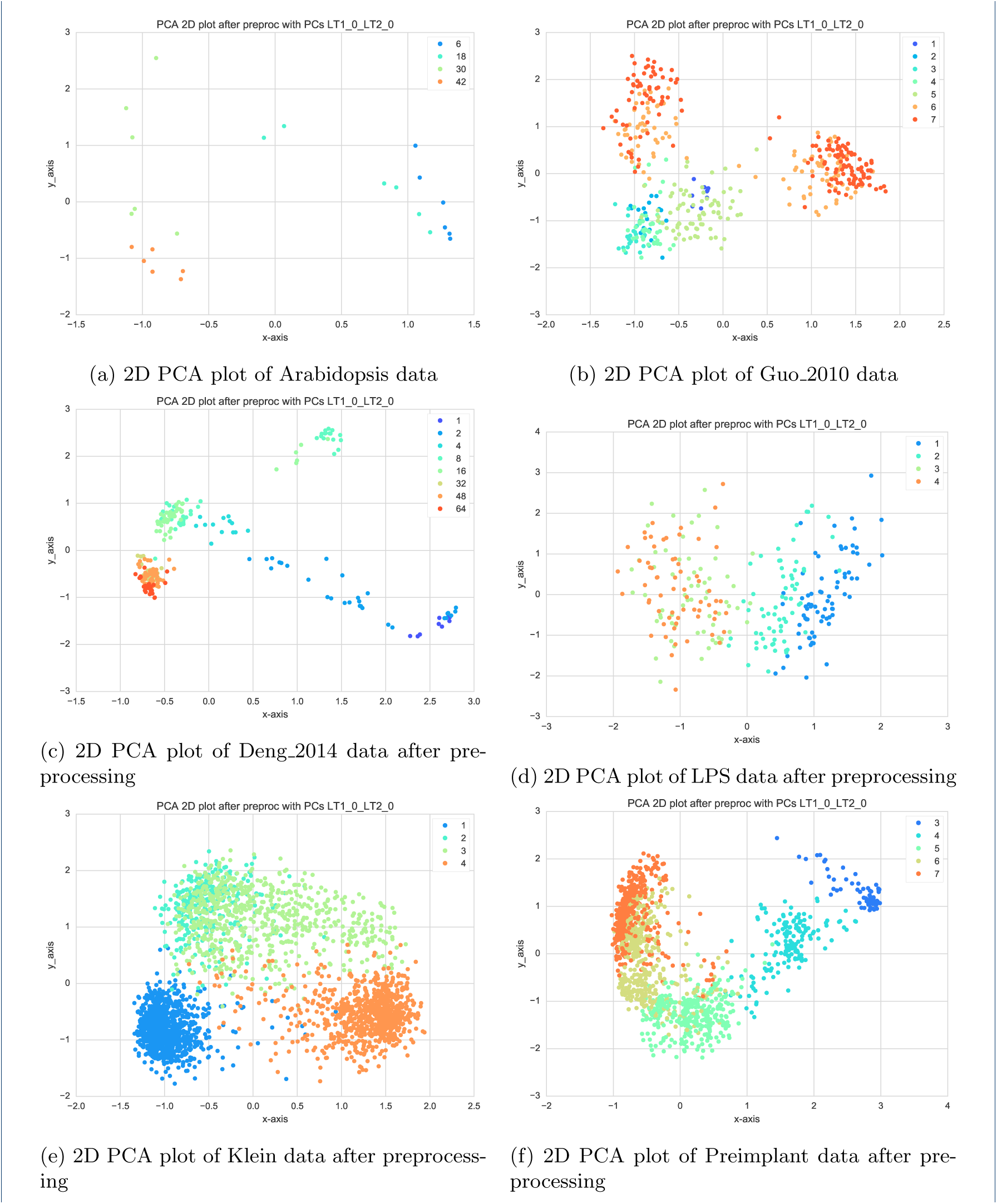
2D PCA plot of Arabidopsis (microarray), Guo_2010 (RT-PCR), Deng_2014 (sc_RNA), Klein (sc_RNA), LPS (sc_RNA) and Preimplant (sc_RNA), the legend shows the various time points present in the datasets

**Table 1:** Six datasets from multiple methods involving microarray, RT-PCR and scRNA-seq are used in the experiments

| Index | Dataset Name | Data type | Original dimension (cells X genes) | Dimension after preprocessing (cells X genes) | Time points | Dataset description | Bifurcations |
| --- | --- | --- | --- | --- | --- | --- | --- |
| 1 | Arabidopsis | microarray | $24 \times 150$ | $24 \times 117$ | 4 | Microarray data of the response of *Arabidopsis thaliana* to infection by necrotrophic fungal pathogen *Botrytis cinerea* | No |
| 2 | Deng 2014 | scRNA-seq | $293 \times 1000$ | $253 \times 994$ | 9 | scRNA-seq embryonic developmental data from mouse preimplantation ranging from zygote to blastocyst | No |
| 3 | Guo 2010 | RT-PCR | $438 \times 48$ | $438 \times 48$ | 7 | RT-PCR data quantifying developmental stages (1-cell to 64-cell blastocyst) from early stage mouse embryo | Yes |
| 4 | Klein | scRNA-seq | $2717 \times 24174$ | $2717 \times 323$ | 4 | Droplet sequencing of mouse embryonic stem cells upon leukemia inhibitory factor (LIF) removal | Yes |
| 5 | LPS | scRNA-seq | $306 \times 27723$ | $268 \times 3910$ | 4 | scRNA-seq bone marrow derived dendritic cell samples after simulating with lipopolysaccharide (LPS) | No |
| 6 | Preimplant embryo | scRNA-seq | $1529 \times 24444$ | $1329 \times 5490$ | 5 | scRNA-seq human preimplantation embryo dataset collected from 88 embryos ranging from day 3 to day 7 | No |

### 2.2 Trajectory Inference

In order to robustly infer single cell differentiation trajectories FORKS assumes the following facts about the data:

1. Differentiation and stimulus response are continuous processes and the data points represent samples on the a continuous manifold.
2. Cells may nevertheless form clusters in low-dimensional space, either due to discrete sampling of the continuous process or because some intermediate cell states are more long-lived than others.
3. Steiner Points represent the centers of such clusters, and cells are distributed normally around the MST edge joining the two Steiner Points.

#### 2.2.1 Data preprocessing

Gene expression levels inferred from scRNA-seq data suffers from high technical variability due to factors such as variation in RNA quality, incomplete cell lysis, variable reagent concentration and variable processing time. Moreover, many genes are detected only infrequently and at low levels. Thus, it is imperative that we discard low quality cells and weakly detected genes in order to achieve a robust outset. After the preprocessing step, FORKS reduces the dimensionality of data using PCA, constructs the Steiner tree and orders the cells by projecting onto the backbone of the tree details of which can be found in section 4. A basic pseudo-time estimation procedure based on MST is described in algorithm 1.

#### Algorithm 1

Basic pseudo-time estimation algorithm

1: **procedure** w=psEuDoTiME(pre-processed data 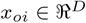, ∀*i* ∈ {1,…, *N*}, reduced dimension *d*, cluster centers *K*)

2: Reduce the dimensionality of the data 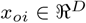 to 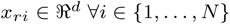

3: Find *K* cluster centers 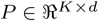

4: Connect cluster centers by a Minimum Spanning Tree

5: Project the data onto tree

6: Allocate a pseudo-time based on the distance from a starting data point

7: **end procedure**

#### 2.2.2 Dimensionality reduction

Even after preprocessing, we are left with thousands of genes, thus leaving us with the task of solving the ordering problem in high dimension. data. However, most of the information presumably lies close to a low dimensional manifold. We therefore reduce the dimensionality of the original data matrix denoted by 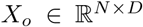 to 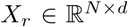, where *N* is the number of cells or data samples, *D* is the number of genes after preprocessing and *d* is the reduced dimension. For this step, we compared the correlation of pseudo-time inferred by the algorithm 1 with the true pseudo-time on several manifold learning/dimensionality reduction algorithms such as PCA [14], ISOMAP [35], Multi-Dimensional Scaling (MDS) [36, 37], Spectral Embedding (SE) [38, 39], Random Forest (RF) [40] and t-SNE [17]. Once the reduced dimensional embedding has been found we cluster the cells.

#### 2.2.3 Spanning tree construction using cluster centers

It is known that similar cell types tends to cluster together. Hence, once a lower dimensional embedding has been found, we find clusters present in the data. These cluster centers are connected via a Minimum Spanning Tree (MST). Finally, the cells are projected onto the MST and distances along the MST from a starting point acts as pseudo-time. We compare various clustering algorithms like k-means, k-medoids, steiner kmedoids (Approximate Euclidean Steiner Minimal Tree (ESMT) algorithm with k-medoids centers as warm start) along side FORKS (steiner kmeans; Approximate ESMT algorithm with k-means centers as warm start) which simultaneously minimizes the clustering objective along with the length of MST joining those centers. These centers are called as Steiner points in case of FORKS.

The effect of various embeddings on average Spearman correlation for various datasets using k-means clustering are shown as box plots in Supplementary Figure 1 (1a-1f). We find that PCA has the highest median correlation on 5 out of 6 datasets. The run times for k-means clustering are also compare the results of whose are found in Supplementary Figure 2 (2a-2f). In this case PCA has the fastest median time for 3 out of 6 datasets.

Supplementary Figures 3 (3a-3b) and 4 (4a-4b) summarizes the above results and displays that PCA has the highest median correlation for all the datasets. It is highly robust with smallest standard deviation of median correlation and is comparable to ISOMAP and MDS in run times.

We then compared the correlations and time for k-medoids clustering. The results are presented, in Supplementary Figures 5 (5a-5f) and 6 (6a-6f) respectively. ISOMAP has highest correlation among all the embeddings for 4 out of 6 datasets.

Supplementary Figures 7 (7a-7b) and 8 (8a-8b) displays that ISOMAP has the highest median correlation for all the datasets.

For FORKS (Steiner tree with k-means warm start), the correlations and run time are shown in Supplementary Figures 9 (9a-9f) and 10 (10a-10f).

Supplementary Figures 11 (11a-11b) and 12 (12a-12b) displays that PCA has the highest median correlation for all the datasets. It is highly robust with smallest standard deviation of median correlation.

Finally, for Steiner tree with k-medoids warm start, the correlations and run times are presented in Supplementary Figures 13 (13a-13f) and 14 (14a-14f).

Supplementary Figures 15 (15a-15b) and 16 (16a-16b) shows that PCA has the highest median correlation for all the datasets while ISOMAP has fastest running time. Although t-SNE displays the most robust behavior it lags considerably in median correlation and runtimes.

We find that PCA has the best average Spearman correlations for k-means clustering, Steiner tree with k-medoids warm start and FORKS (Steiner tree with k-means warm start). ISOMAP has the best average Spearman correlation for k-medoids clustering. The average Spearman correlation for PCA as an embedding is much higher than that of ISOMAP which becomes the basis of choosing PCA as the choice of manifold learning.

Finally, we compared several clustering algorithms on different datasets and embeddings. Supplementary Figures 21 (21a -21d), shows that k-means and FORKS have similar average Spearman correlations, but FORKS is more robust compared to k-means as it has a smaller median standard deviation (0.058 compared to 0.065) across various algorithms and datasets.

These results provide empirical evidence for using PCA as dimensionality reduction technique and Apporximate ESMT as the method of choice to find the cluster centers in FORKS. In the next subsection 2.3, we compare FORKS to 8 other state-of-the-art algorithms.

### 2.3 Comparison with other algorithms

We compared FORKS with eight other state-of-the-art algorithms namely DPT, GPfates, kmeans-R (used in [25]), Monocle2, SCUBA, SLICER and waterfall. The description of the algorithms used, viz., the dimensionality reduction techniques, trajectory modeling framework, scalability and their ease of use is mentioned in Tables 2 and 3. As a preprocessing step we use pQ normalization [49] for scRNA-seq datasets and data spherization (zero mean and unit variance transformation) for microarray and RTPCR datasets. The data was then divided using stratified k-fold technique used in scikit [50] such that each fold contains similar distribution of true cell times as original data. The value of *k* varies with dataset due to difference in number of cells sequenced. Keeping the folds and the dataset same, all the algorithms were benchmarked. We calculated the Spearman correlation [51] between the pseudo-time computed by algorithm and the true time for each fold. Supplementary Table 1 shows the (mean ± standard deviation) Spearman correlation for each of the algorithm. Among all the benchmarked algorithms, Monocle2, SCUBA, TSCAN, kmeans-R and waterfall do not require information about the ’starting cell’ of the pseudo-time. Hence, we ran all these algorithm for a forward pass and a backward pass from their internal random starting point. For providing a more competitive environment to FORKS, we took the absolute values of correlation for these algorithms. ^[1]^

**Table 2:**
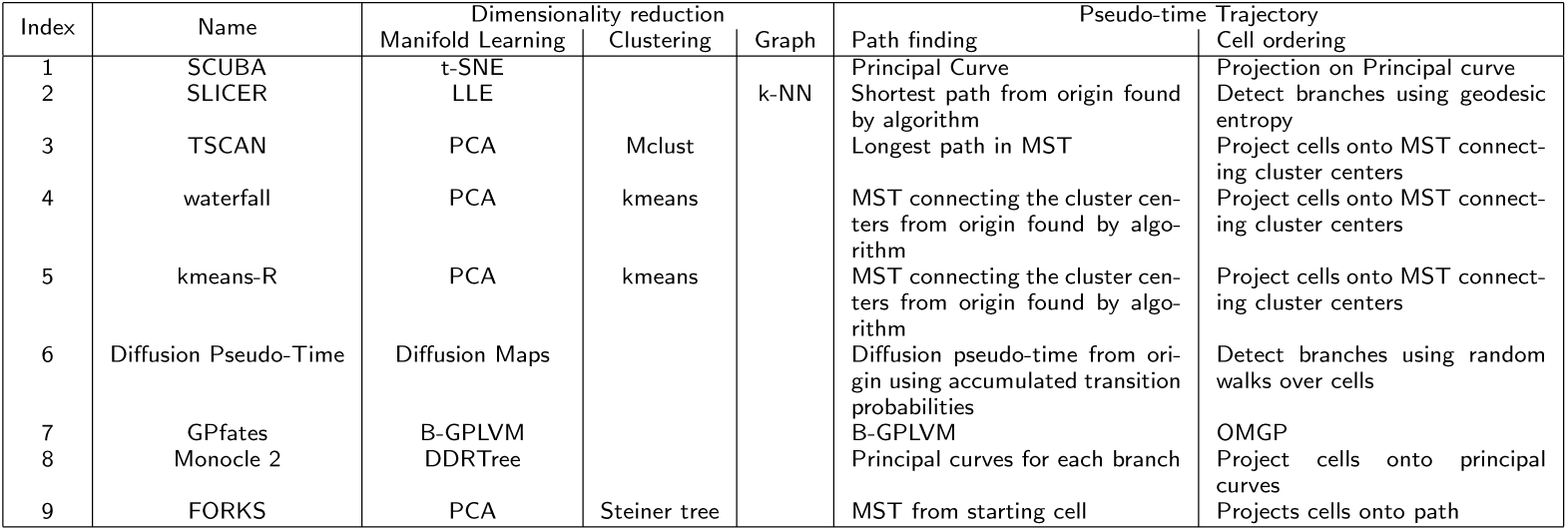
Pseudo-temporal ordering methods have many commonalities in their framework. As with all the methods, the first step being dimensionality reduction, then clustering or graph based analysis helps to easily model the trajectory, various methods are then utilized to find the ordering.

**Table 3:**
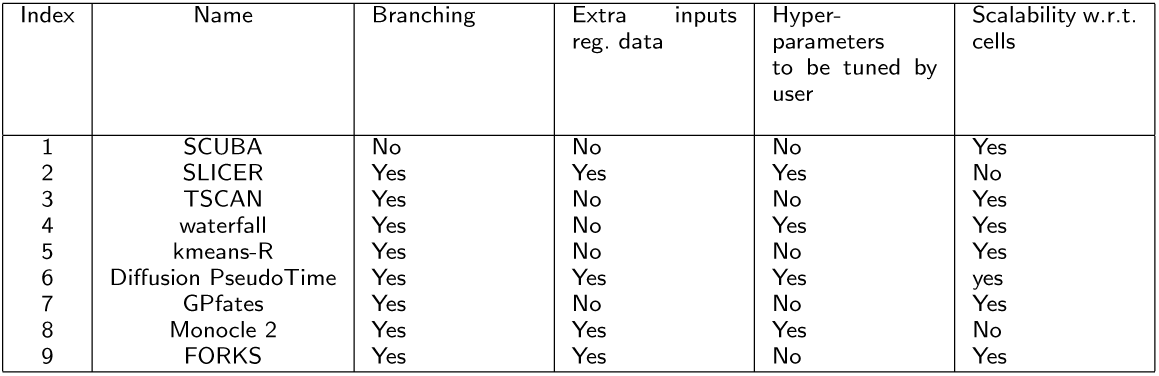
Table describes the scalability and amount of user interference required

Figure 2 (2a-2f) shows the box plots of Spearman correlation for various folds on different datasets. Figure 2 show that FORKS outperform other algorithms in terms of Spearman correlations with known cell times in which median Spearman correlation is greater than 0.9 in 5 out of 6 datasets.

**Figure 2:**
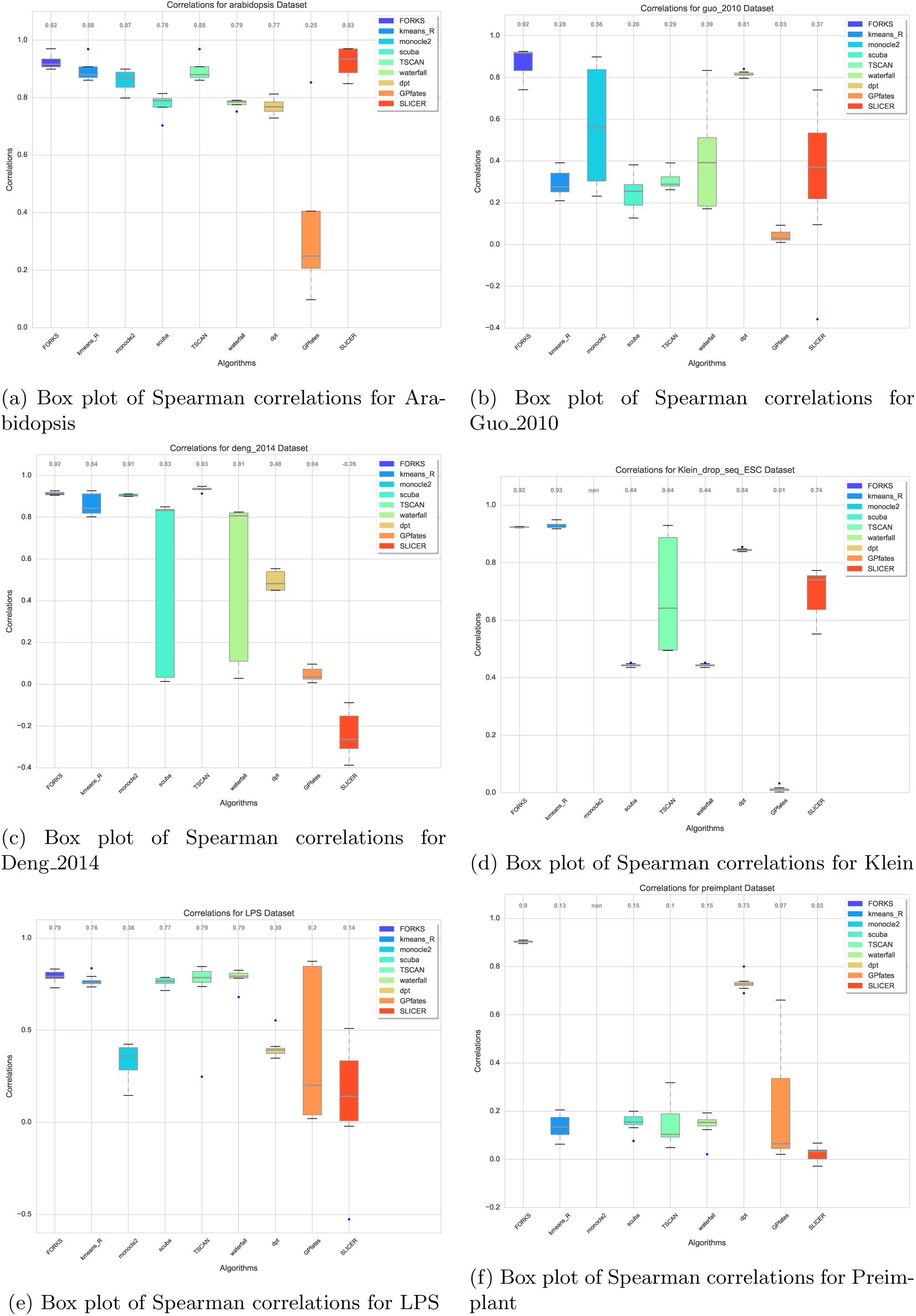
Box plots showing the correlations with given cell times for various algorithms for Arabidopsis (nfolds=4), Guo_2010 (nfolds=8), Deng_2014 (nfolds=5), Klein (nfolds=10), LPS (nfolds=8) and Preimplant (nfolds=10) datasets, the legend shows the various algorithms being compared. Values at the top of each figures are the median values

Run times of various algorithms were also compared. Figures 3 (3a-3f) displays the box plots of run times in seconds, for various algorithm. Although it may not be fair to compare the run times of various algorithms as they are written in multiple programming languages. FORKS is completely coded in Python [52], and uses NumPy [53], SciPy [54] and Pandas [55] internally. FORKS has a higher run times than algorithms whose back-end is written in C or C++ for certain datasets. Still considering these factors, FORKS is competitive in run times to other algorithms and in multiple instances where the dataset sizes are large it is faster than most. Figure 5 (5a-5b) shows the mean and standard deviation run times of all algorithms. We find that FORKS is third fastest in terms of run time and fourth in terms of robustness with respect to run times.

**Figure 3:**
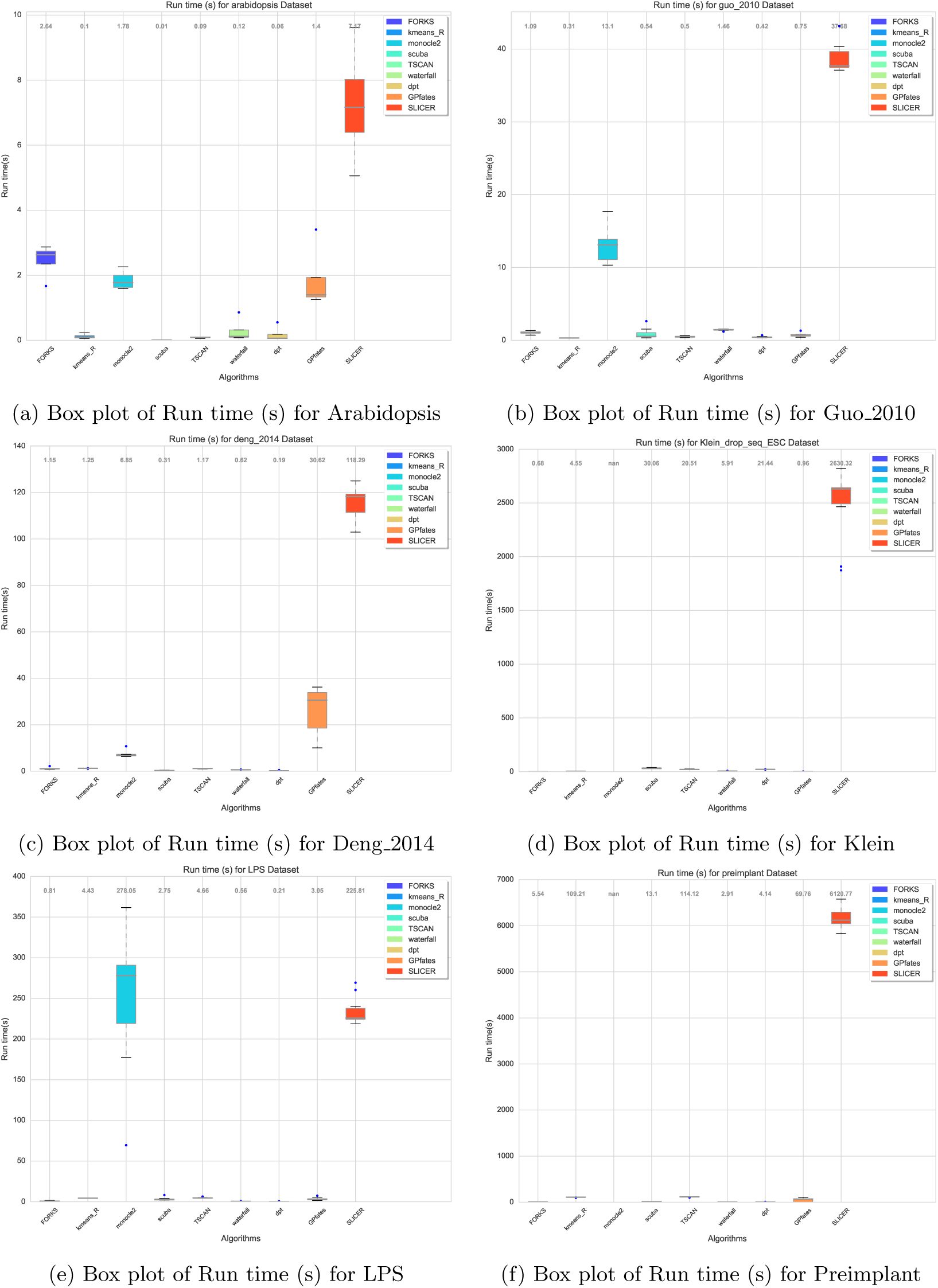
Box plots showing the run times (s) with given cell times for various algorithms for Arabidopsis (nfolds=4), Guo_2010 (nfolds=8), Deng_2014 (nfolds=5), Klein (nfolds=10), LPS (nfolds=8) and Preimplant (nfolds=10) datasets, the legend shows the various algorithms being compared. Values at the top of each figures are the median values

In order to check the accuracy and robustness of each method, we computed the mean and standard deviation of mean Spearman correlations for multiple instances of re-sampling of datasets which we call as data folds. Knowing the true cell times, we used stratified k-fold techniques to generate folds for each dataset. For an accurate and robust algorithm, its mean correlation across folds should be close to 1 and its standard deviation should be close to 0. The results are shown in Figure 4(4a-4b). FORKS has the highest median of the mean correlation among all the algorithms. Figure 4 indicate that FORKS has 10% higher median than its nearest competitor kmeans-R which is at 0.81. Using such an algorithm whose median correlation lies near 0.9 can faithfully recover the true trajectory for most of the datasets. FORKS also exhibits robust behavior across multiple datasets as the mean standard deviation achieved by the algorithm is the smallest at 0.02.

**Figure 4:**
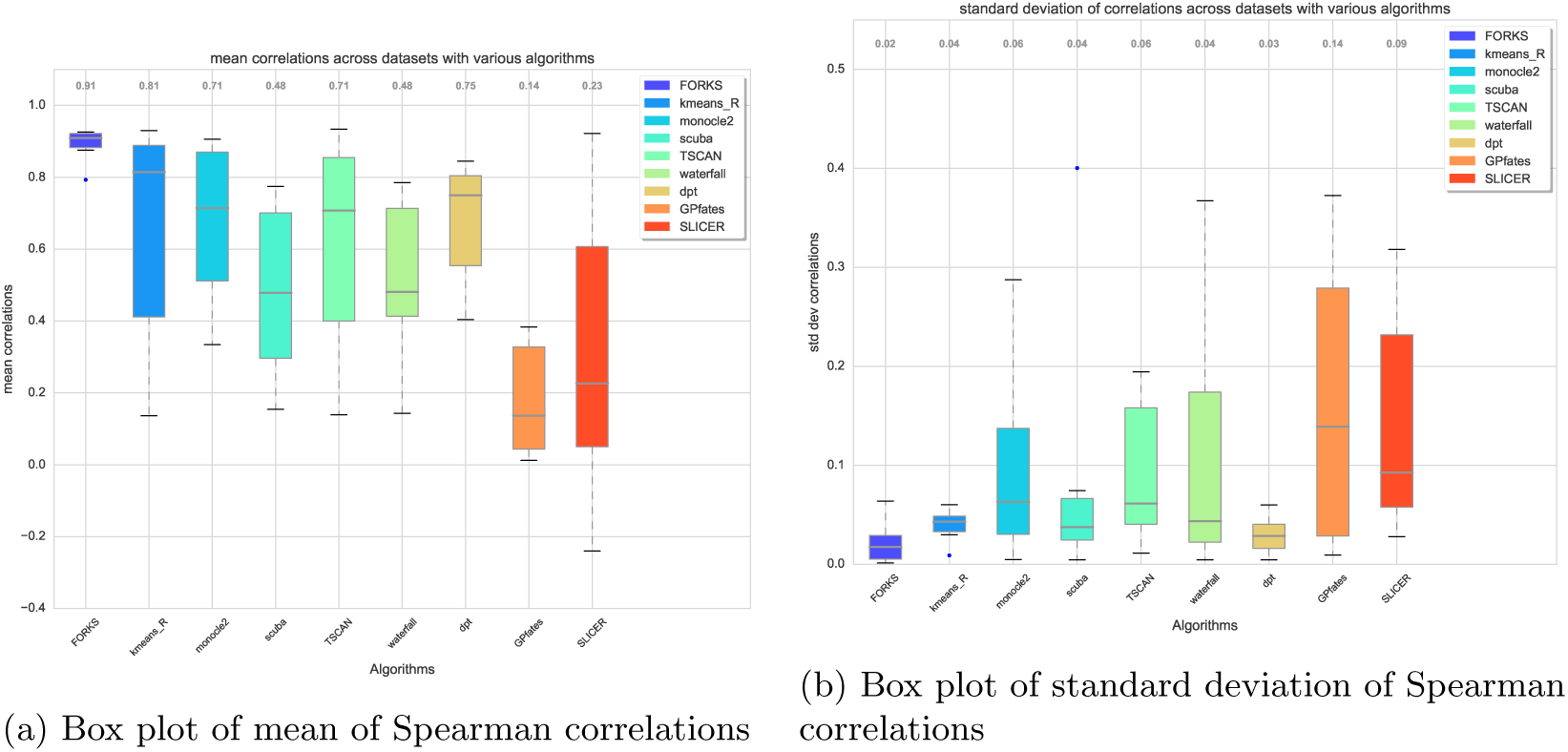
Box plots of means and standard deviation of Spearman correlations among all the datasets shows highly accurate and robust behavior of FORKS to change in folds and datasets. Values at the top of each figures are the median values

**Figure 5:**
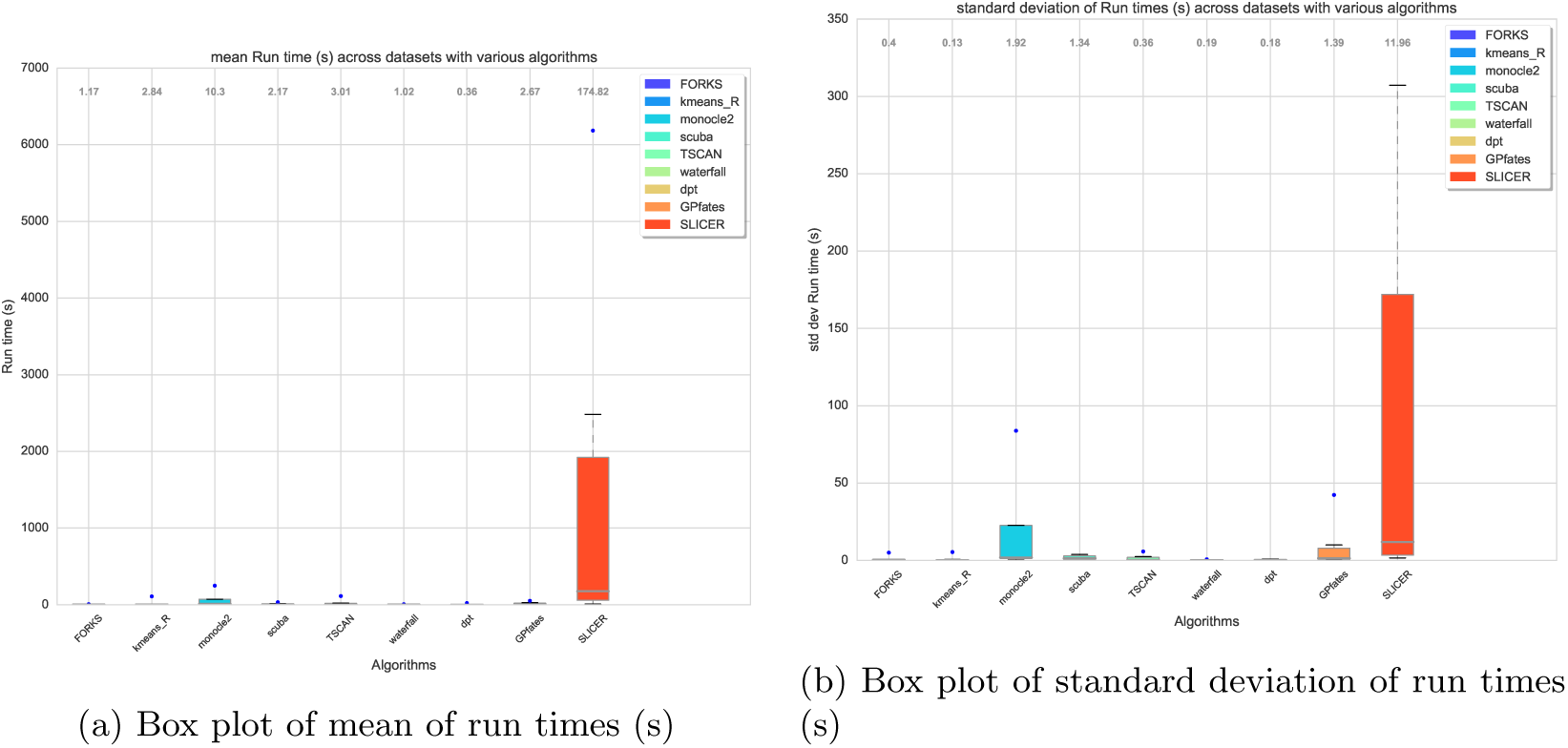
Box plots of means and standard deviation of run times among all the datasets shows FORKS is highly competitive. Values at the top of each figures are the median values

## 3 Discussion

We developed FORKS (Finding Orderings Robustly using k-means and Steiner trees), a method that infers cellular trajectories from various gene expression datasets. Having the highest average Spearman correlation and smallest mean standard deviation of Spearman correlation with known cell times compared to other algorithms, FORKS is able to robustly and accurately discover branching trajectories if present in the data. One of its biggest strength being the robustness of the default parameter setting for multiple datasets, an advantage which is not possessed by its competitor algorithms. FORKS uses PCA as the dimensionality reduction technique and infers the trajectory via projection on Steiner tree. The rationale behind using PCA as the manifold learning algorithm is described in subsection 3.1.

### 3.1 Using PCA as manifold learning algorithm of choice

Manifold learning is a crucial step involved in pseudo-temporal ordering. For this step, we compared several manifold learning/dimensionality reduction algorithms such as PCA [14], ISOMAP [35], Multi-Dimensional Scaling (MDS) [36, 37], Spectral Embedding (SE) [38, 39], Random Forest (RF) [40] and t-SNE [17]. Contrary to prevailing biases towards using non-linear dimensionality reduction methods like Diffusion Maps [22], our experiments demonstrate the preeminence of PCA over these techniques. Advantages of PCA includes its computation speed, scalability and the fact that the number of dimensions to project the data on can be calculated using the energy or the variance captured in each component. We found that other methods to find the reduced dimensional embeddings are highly sensitive to other parameters such as kernel width as in case of Gaussian kernel for kernel PCA [41] or Diffusion Maps, perplexity in case of t-SNE and number of nearest neighbors in case of ISOMAP, Locally Linear Embedding (LLE) or any other method based on k-nearest neighbor graph.

### 3.2 Using Steiner-Tree with k-means as clustering algorithm of choice

The Euclidean Steiner Minimal Trees (ESMT) problem, as it is called, finds the shortest tree connecting *N* data points with *K* Steiner points, where *K* ≪ *N*. ESMTs have been successfully applied to the areas of very large integrated circuits (VLSI), printed circuit boards, in the field of agriculture, telephony and in optimizing the building layouts. Application of ESMTs in the domain of genomics is the bridge we try to fill through this work. (ESMT) finds the Steiner points or cluster centers such that the total length of spanning tree formed by joining the Steiner points/cluster centers is minimum. In many cases its solution is very similar to the solution of k-means but in case of noisy data, it finds robust solutions as compared to k-means. Steiner Minimal Trees (SMTs) are hard to find as the number of Steiner points are not known a priori and for a given number of Steiner points, one has to determine the correct topology with respect to the other data points. It has been shown that computation of SMTs is NP-complete [42]. A work around this problem is to set the number of Steiner points and then solve an optimization problem to determine the approximate location of Steiner points. The algorithm proposed in this paper is termed as Approximate Euclidean Steiner Minimal Trees abbreviated as approximate ESMT. ESMT problem has a strong connection with k-means and Minimum Spanning Tree (MST). We propose a gradient based optimization approach to solve the NP-complete problem. Owing to non-convex nature of optimization problem the algorithm may get stuck in a local minima. Once the MST joining the cluster centers is found, we project the cells onto the tree backbone or MST joining the cluster centers. FORKS uses a ’starting cell’ to commence the ordering. It finds the cluster center closest to the ’starting cell’ and start the ordering from that center which we call as root of the tree. A starting cell is necessary because a completely unsupervised method may choose any of the points of MST to start the ordering, which might result in an incorrect ordering. Distances along the tree backbone starting from the root Steiner point serves as a proxy for pseudo-time.

Using the Steiner points to construct the MST, our algorithm can find bifurcations using small linear segments. When cross validated with multiple stratified folds of the datasets, we find that FORKS is superior to the other eight competing algorithms in terms of highest Spearman correlation with true pseudo-time. FORKS is highly robust to change in datasets and folds which is attributed to its smallest standard deviation of Spearman correlation on multiple datasets and their respective partitions. It requires *O*(*NK*) memory for storage of distance matrix hence can be scaled to large datasets. The name FORKS is an abbreviation for Finding Ordering Robustly using k-means and Steiner tree as it aptly finds a robust pseudo-temporal trajectory with the combination of k-means and Steiner Tree.

### 3.3 Conclusion

In this paper we presented FORKS, an algorithm that finds a robust pseudo-temporal ordering of cells with no tuning of hyper-parameters. We compared FORKS with several state-of-the-art algorithms and showed its supremacy over them. We demonstrated that FORKS can be used with any kind of gene expression dataset till the dataset is in a tabular numeric form of cells × genes. We empirically proved our claim that the task of pseudotemporal ordering can also be solved effectively using linear embedding contrary to the general notion prevalent in the bio-informatics community. Finally, we incorporated the ideas from VLSI domain in the form of steiner tree and solved its approximate version using gradient descent to device FORKS.

With ever increasing dataset size and quality, methods that scale up will have a certain edge over algorithms that do not. FORKS was created with such an ideology. Being closely related to k-means and using PCA as dimensionality reduction technique both of which can be solved stochastically, FORKS can be scaled up to large datasets. There is always scope of improving the current algorithm. Gene selection is an important step in any downstream analysis. For the purposes of this paper, we have used only the highly expressed and varying genes. For the problem at hand, using the marker genes can certainly improve the ordering. Also, to make the run times more smaller, the code can be cythonized or be completely coded in hardware friendly languages like C or C++. Also, one can try other clustering algorithms like Gaussian Mixture models (GMMs) [56], DBSCAN [57] and kernel-kmeans [58] to find the cluster centers and measure its performance.

## 4 Online Methods

### 4.1 Data Preprocessing

Dissimilitudes present in single cell data presents us with the challenge of adopting the right processing for each kind of dataset. We first highlight the common steps followed for all the datasets. The genes and cells containing all zero elements along with the duplicated genes and cells present in the data were removed. Since the marker gene information is not available in all the datasets and in the light of making the algorithm largely free from human intervention, we incorporated a gene selection step. Only a few genes who are expressed faithfully are responsible for the trajectory inference in the cells. We find the median and standard deviation of all the non-zero genes present in the data, by forming a one dimensional array. We discard low quality genes whose individual median gene expression in the expressed cells is below a certain multiple of the overall median.

For the single-cell RNA sequencing datasets namely, (Klein [46], LPS [47], Deng_2014 [44], Preimplant [48]), we performed pseudo-counted Quantile (pQ) normalization [49]. It is well known that single cell RNA-sequencing datasets have inherent technical variability and high biological noise [59, 60, 61]. pQ normalization is a recently proposed technique build upon Quantile (Q) normalization [62] that reduces the technical bias, which is empirically found to be directly proportional to the number of detected genes [63]. pQ normalization homogenizes the expression of all genes below a fixed rank in each cell. Finally, for non-single cell RNA seq datasets (Guo_2010 [45] and Arabidopsis [43]), we found that sphering the dataset improves the pseudo-time estimation. However pQ normalization degrades the performance.

### 4.2 Overview of FORKS algorithm

It is known that only a few interactions are responsible for a particular problem like cell cycle or differentiation. In case of scRNA-seq, lowly expressed genes contribute largely to technical bias. Hence, it is imperative to reduce the dimension to capture the most meaningful interactions.

#### Dimensionality reduction

The algorithm begins by dimensionality reduction. Reducing the dimension, diminishes the complexity of problem at hand and reduces the noise present in higher dimensions. We use Principal Component Analysis (PCA) [14] to reduce the dimension of the data. PCA offers a graceful solution to problem of selecting the required number of dimension, by using the energy content in each of the principal components (PCs). We select the PCs such that the total energy content is at least 90%. There are other approaches to do so, for example, by eyeballing the curve to get the knee point. However, due to the lack of mathematical justification, this approach was discarded. PCA is a linear manifold learning technique many other pseudo-temporal ordering methods use various non-linear manifold learning techniques, and pay the cost of tuning multiple hyper-parameters, thereby increasing the complexity of training. If the parameters are not chosen correctly, one might end up learning erroneous manifold thereby deducing incorrect pseudo-time. PCA in this regards provides an excellent solution.

#### Finding number of clusters

Initial release of Monocle used minimum spanning tree (MST) based on the complete data and then found the longest path present in the MST. One major issue with this approach was the fact that MST on such a large dataset is not stable. Clustering the cells first and choosing cluster centers as the proxy for data is one way to mitigate problem. Once the dimension is reduced, next we choose the number of clusters that might be present in the data using Silhouette scores [64]. When the data labels are not known, the cluster evaluation should be performed using the model itself. The Silhouette Coefficient is composed of two scores for each sample:

1. *a*: The mean distance between a sample and all other points in the same class.
2. *b*: The mean distance between a sample and all other points in the next nearest cluster

The Silhouette Coefficient (s) per sample is calculated as:

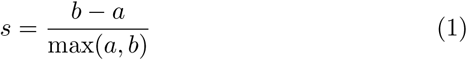

The Silhouette Coefficient for a set of samples is given as the mean of the Silhouette Coefficient for each sample. The best value is 1 and the worst value is −1. Values near 0 indicate overlapping clusters. Negative values generally indicate that a sample has been assigned to the wrong cluster. We calculate the silhouette scores for the cluster centers ranging from 4 to 10. The number of cluster centers which gives the best average Silhouette Coefficient are selected.ESMTLet the number of cluster centers/steiner points be *K* ≪ *N*. The cluster centers/steiner points are denoted by 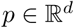 and data points by 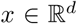. Let 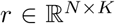 be the matrix denoting the cluster center data point is assigned to. The problem of finding the Euclidean Steiner Minimal Tree (ESMT) can be written as:

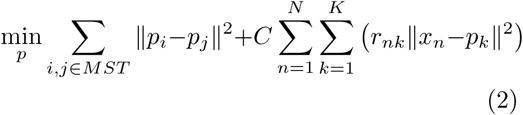

where, MST denotes the minimum spanning tree formed by steiner points.

The ESMT problem given as eq. (2) is non-convex as the second part of the objective function is the same as that of k-means, which is not a convex function [65]. Hence, optimization leads to a local minima. We use gradient descent to update the steiner points. As mentioned in the section 2, finding the Steiner tree is a NP-hard problem. We use approximate ESMT where we fix the number of clusters a priori. The initial number of cluster centers are calculated using Silhouette method and their initial values are computed using the solution of k-means. The hyper-parameter *C* is set to 1 in all our experiments. FORKS, which is mathematically written as eq. (2) can be interpreted as finding the steiner points such that the total length of minimum spanning tree formed joining the steiner points is minimum. We run the algorithm for 100 epochs keeping track of the best cost function values. The values of steiner points for which the cost function is minimum is returned.

#### Ordering

We divide the points belonging to each cluster and then use the methodology mentioned in TSCAN [25] for pseudo-temporal ordering. We first select a ”starting cell” based on user input. For our experiments in order to demonstrate a proof of concept, we select the first cell belonging to the initial time as the starting cell.

Selecting a starting cell is important and this is where we have an edge over TSCAN or other methods which do not have a provision of selecting a starting cell. The proposed method of finding the longest path [25] in the MST can lead to erroneous results as it can be seen through the Figure 2 where TSCAN does not preform well. For bifurcating trajectories as in case of Figure 1b, where initial stage cell population may lie in the between the later stages of cell population, selecting the starting cell for ordering as one of the ends of the longest path is incorrect.

The ordering begins from the steiner point *k*_1_ closest to the starting cell. Cells belonging to steiner point *k*_1_ are assigned to edge *k*_1_ − *k*_2_. For, the intermediate steiner points *k_i_*, (*i* = 2, …, *K* − 1), the cells belonging to cluster *k_i_* are divided into *t* parts where, *t* is the number of immediate neighbors of *k_i_* in the MST. Once the assignment of cells is finished for each edge *e_j_* ∈ *MST* in MST, the cells are projected on their respective edges. The cell closest to *k*_1_ is assigned a pseudo-time of 0. All the other cells are assigned values equal to the order of sorted projection value onto the edge *e_i_* added with the last value of its preceding edge.

## Code

The code for FORKS along with three of the datasets used in the paper can be found at the following github repository https://github.com/macsharma/FORKS.

## Competing interests

The authors declare that they have no competing interests.

## Author’s contributions

## Acknowledgements

Authors would like to acknowledge Tim Xiaoming for his feedback on scRNA-seq datasets and improvements to FORKS.

## Footnotes

[1] All the tests were performed on a machine with Intel core-I5 2nd Gen processor with 6GB DDR2 RAM

